# Dithering suppresses half-harmonic neural synchronisation to photic stimulation in humans

**DOI:** 10.1101/2025.10.03.680216

**Authors:** Benoit Duchet, Samini Subramaniam, Alexander Greenway, Shenghong He, Nicholas Shackle, Alek Pogosyan, Timothy Denison, Andrew Sharott, Huiling Tan, Rafal Bogacz

**Affiliations:** MRC Brain Network Dynamics Unit, Nuffield Department of Clinical Neuroscience, University of Oxford, Oxford, United Kingdom; University of Oxford, Oxford, United Kingdom; Queen Mary University of London, London, United Kingdom; Institute of Biomedical Engineering, Department of Engineering Sciences, University of Oxford, Oxford, United Kingdom

## Abstract

While entraining neural rhythms using brain stimulation has been suggested as a therapeutic mechanism to normalise brain activity in conditions such as depression, chronic pain, or Alzheimer’s disease, periodic stimulation can also inadvertently entrain brain rhythms at sub- and superharmonics of the stimulation frequency, which could lead to deleterious effects. Slightly jittering stimulation pulses (called “dithering”) was previously proposed on the basis of mathematical modelling to selectively entrain a target neural rhythm while avoiding harmonic entrainment. In this study, we investigated the potential of dithering in humans. Using photic stimulation (light flicker) and EEG recordings in healthy participants, we showed that dithering suppresses half-harmonic synchronisation relative to perfectly periodic flicker, and more so than synchronisation at the stimulation frequency. This was also the case for a periodic condition with reduced stimulation amplitude, as predicted by theory. Furthermore, we demonstrated using synthetic data and modelling that the half-harmonic responses observed in participants cannot be explained by the superposition of evoked responses (even when modulated at the half-harmonic frequency), and are better matched by a minimal oscillator model. Our findings are consistent with half-harmonic EEG synchronisation in response to photic stimulation predominantly reflecting half-harmonic entrainment rather than the summation of evoked responses, and with dithering being an effective strategy to suppress subharmonic entrainment.

## 1 Introduction

Entraining neural rhythms using brain stimulation has been suggested as a new therapeutic mechanism to normalise brain activity. For example, emerging approaches target the alpha band (8-12 Hz) using transcranial or sensory stimulation in individuals with depression or chronic pain [1, 2, 3, 4]. Gamma band (30-100 Hz) entrainment also hold promises, using sensory stimulation in Alzheimer’s disease [5, 6, 7, 8], and transcranial alternating current stimulation as well as deep brain stimulation (DBS) in Parkinson’s disease (PD) [9, 10].

However, periodic stimulation can also entrain neuronal rhythms at subharmonics and superharmonics of the stimulation frequency, which could lead to harmful effects. For example, finely-tuned gamma oscillations can be entrained at half the frequency of DBS in patients with PD [11, 12, 13, 14, 15] and dystonia [16]. This subharmonic entrainment was initially thought to promote debilitating involuntary movements known as dyskinesia [11, 12]. While recent studies have uncovered a more complex relationship between this subharmonic entrainment and dyskinesia [17, 15], they support the general principle that unintended entrainment can functionally disconnect neural oscillations, which could in some cases lead to unintended behavioral manifestations. Following this principle, DBS frequency was set to avoid subharmonic entrainment of rhythms associated with epileptic seizures in a canine with epilepsy [18].

To selectively entrain a target neural rhythm while avoiding potential harmful effects from sub- and superharmonic entrainment, a stimulation approach called “dithering” has been proposed [19]. In its simplest form, dithering involves jittering stimulation pulses, such that the duration of each inter-pulse interval varies slightly from the mean stimulation period. Since sub- and superharmonic entrainment are less stable than entrainment at the mean stimulation frequency, a level of dithering can be found that preserves entrainment at the target frequency while suppressing sub- and superharmonic entrainment (Fig 1). The ability of dithered stimulation to achieve selective entrainment was established theoretically and verified computationally in networks of coupled neural oscillators [19], but has not yet been tested experimentally.

**Figure 1:**
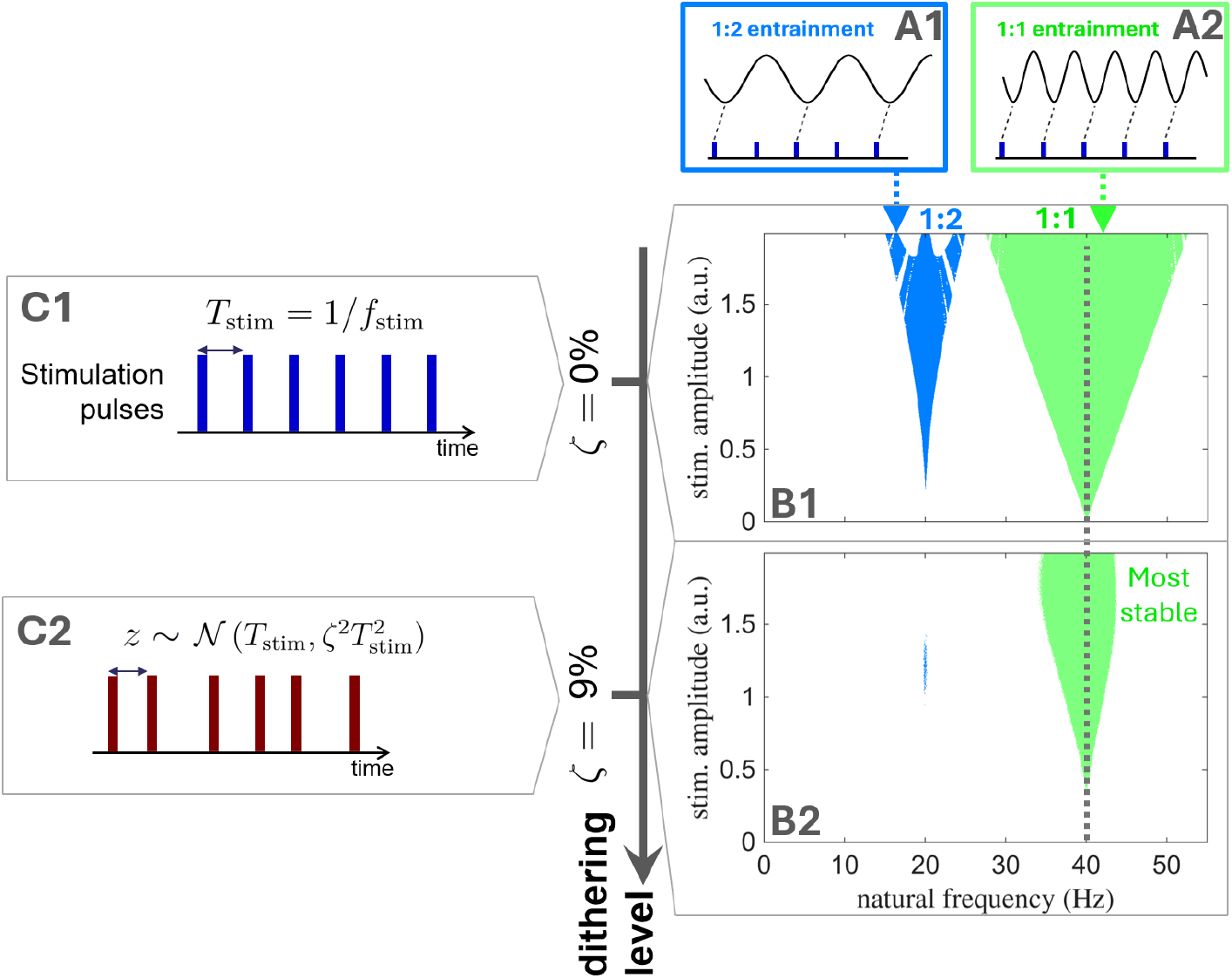
Predicted effect of dithering on subharmonic entrainment. When stimulation is perfectly periodic as depicted in **C1** (*f*_stim_ denotes the stimulation frequency), neural oscillators may be entrained at the stimulation frequency but also at the half-harmonic of the stimulation frequency (as well as superharmonics and other subharmonic ratios, not shown here). Corresponding entrainment regions (called Arnold tongues) are represented in **B1** for uncoupled neural oscillators modelled in [19]. Stimulation is provided at 40 Hz (grey dashed line), with stimulation amplitude shown on the vertical axis, and the natural frequency of oscillators on the horizontal axis. Entrainment to the stimulation frequency (1:1 entrainment) is observed in the green region, while half-harmonic entrainment (1:2 entrainment) is observed in the blue region. Schematics representing both types of entrainment are presented in **A1-2**. With dithering, stimulation pulses are slightly jittered (**C2**), and past a certain dithering level (level of noise in the stimulation period, denoted by *ζ*), only the 1:1 Arnold tongue subsists (green tongue in **B2**). Figure adapted from [19].

Here, we test the ability of dithering to modulate subharmonic synchronisation in healthy humans using photic stimulation (light flicker) and electroencephalography (EEG). The ongoing debate on whether photic stimulation entrains neural activity at the stimulation frequency (with neural oscillators synchronising to the stimulation frequency) or simply evokes time-locked neural responses to each individual flash [20, 21, 22, 23, 24] precludes us from concluding on the impact of dithering on entrainment at the stimulation frequency and its superharmonics. Instead we consider subharmonic responses to photic stimulation, which have been reported in numerous human studies [25, 26, 27, 28, 29, 6]. We show that these subharmonic responses can be suppressed by dithered stimulation, and present evidence that these subharmonic responses are not consistent with evoked responses and likely reflect subharmonic entrainment.

## 2 Results

Since there is individual variability in which stimulation frequencies lead to subharmonic responses to photic stimulation, we first identified for each participant the stimulation frequency leading to the largest power response at the half-harmonic (denoted *f*_max 1:2_) using a frequency sweep from 15 to 43 Hz (Fig 2A1). For the included datasets, the mean *f*_max 1:2_ was 32.4 ± 6.7 Hz (individual values reported in Table B in the supplementary material, also see example response to stimulation frequency sweep in Fig S.1D in the supplementary material).

**Figure 2:**
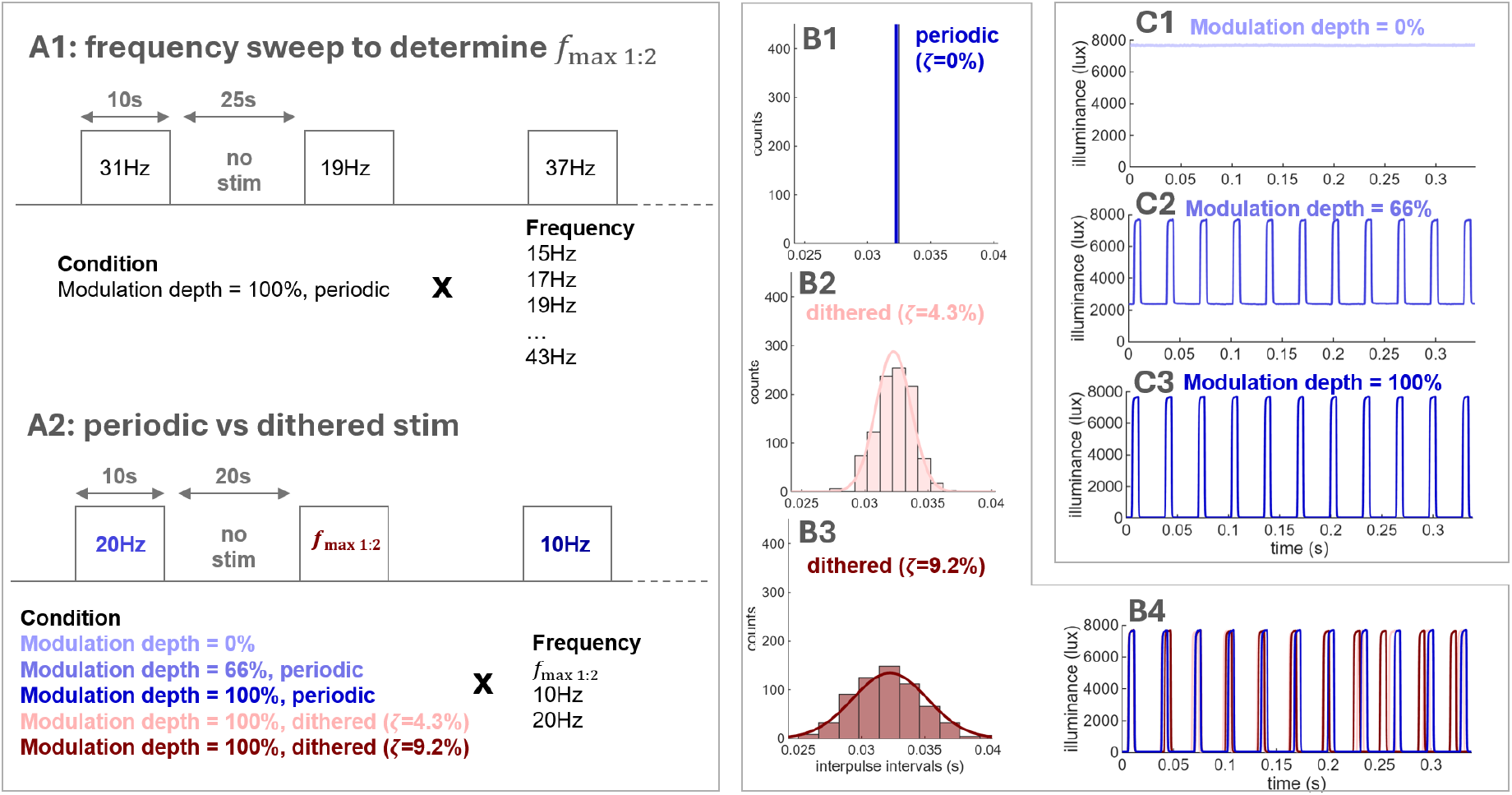
Experimental paradigm and stimulation conditions. **A1**: Schematic of the frequency sweep protocol used to determine *f*_max 1:2_. **A2**: Schematic of the protocol used to compare dithered stimulation to periodic stimulation. **B**: Distribution of inter-pulse intervals for the periodic and dithered conditions (B1-B3). The dithering level *ζ* scales the standard deviation of the inter-pulse interval distribution. B4 shows example stimulation trains for the three conditions. **C**: Stimulation trains with different modulation depths. The modulation depth inversely scales the “off” part of the stimulation pulse. A modulation depth of 0% corresponds to continuous illumination (control condition, C1). Examples in panels B-C are from one participant with a stimulation frequency of 31 Hz.

We next assessed the effect of dithering on half-harmonic synchronisation (Fig 2A2), and used synthetic data and modelling to characterise the nature of this half-harmonic synchronisation. Dithering was implemented by sampling the time interval from one stimulation pulse to the next from a normal distribution 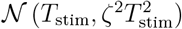, where *T*_stim_ = 1*/f*_stim_, with *f*_stim_ the target stimulation frequency, and *ζ* the dithering level (Fig 2B). Following previous model predictions [19] (Fig 1B2), we considered *ζ* = 9.2% (which is sufficient for the 1:2 Arnold tongue to disappear), *ζ* = 4.3% (an intermediate level), and *ζ* = 0% (the periodic case).

Since theory also predicts that sufficiently reducing stimulation amplitude will suppress 1:2 entrainment (the system will leave the 1:2 Arnold tongue, see Fig 1B1), a periodic condition with a reduced modulation depth of 66% was also included (Fig 2C2). Finally, we included a control condition (modulation depth of 0%, i.e. continuous illumination without flicker, Fig 2C1).

After data recording, we excluded six datasets from further analysis due to the lack of subharmonic response to periodic stimulation or too many rejected trials (see detailed rejection criteria in Section 4.4), resulting in 10 participants included for analysis.

### 2.1 Dithering suppresses half-harmonic responses

Dithering reduced the power of the half-harmonic response to photic stimulation compared to periodic stimulation at *f*_max 1:2_. The power response of a representative participant is presented in Fig 3A and shows a greater reduction in power at the half-harmonic for *ζ* = 9.2% than for *ζ* = 4.3%. Because of between-participant differences in baseline power and maximum power response, we considered for group-level analysis the “normalised power above control”, defined as the difference between the power response for the condition of interest and the control condition, normalised by the difference between the power response of the periodic condition with full modulation depth (which elicits the strongest response) and the control condition. As seen in Fig 3B, both dithering levels (as well as the reduced modulation depth condition) significantly reduced the normalised power above control at the half-harmonic (p < 0.001 in all three cases, one-tailed). For both dithering levels, there was a trend in dithering reducing power at the half-harmonic of stimulation more than at the stimulation frequency (comparison of ratios relative to the periodic condition with full modulation depth at 1:1 and 1:2, p=0.08 in both cases, one-tailed). The reduced modulation depth condition significantly decreased power at the half-harmonic of stimulation more than at the stimulation frequency (comparison of ratios relative to the periodic condition with full modulation depth at 1:1 and 1:2, p < 0.001, one-tailed). Since the power of the response is a coarse measure of synchronisation with stimulation (increases in power do not always correlate with increases in synchronisation [30, 31]), we assessed synchronisation in a more specific manner using the phase-locking value (PLV).

**Figure 3:**
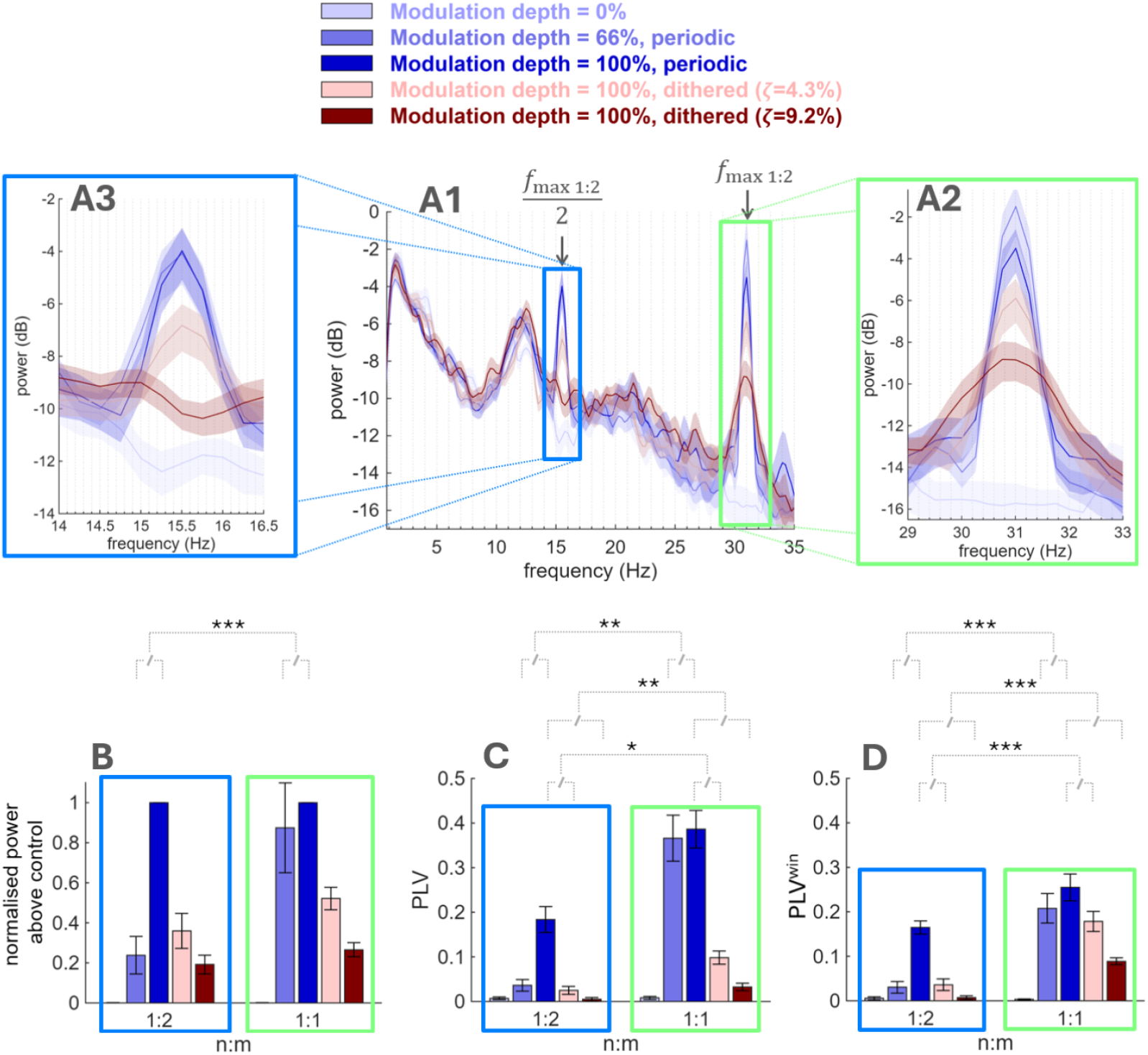
Dithering suppresses half-harmonic synchronisation. **A1**: Power spectrum of participant 14 (averaged over trials and EEG channels) in response to periodic (shades of blue) and dithered (shades of red) photic stimulation at *f*_max 1:2_ = 31 Hz. The lightest shade of blue corresponds to a control condition (continuous illumination, no flicker). **A2** and **A3** are zoomed-in inserts centered on the stimulation frequency and its half-harmonic, respectively. **B**: normalised power above control at the stimulation frequency (1:1) and its half-harmonic (1:2) at the group level. **C**: PLV at the stimulation frequency (1:1) and its half-harmonic (1:2) at the group level. **D**: Windowed PLV at the stimulation frequency (1:1) and its half-harmonic (1:2) at the group level. In panels B-D, n = 10, and significance is only indicated between ratios of conditions relative to the periodic condition with full modulation depth to avoid clutter. In panels C-D, the control condition has a PLV close to zero as expected (the noise contribution is removed in our PLV measure). All panels share the same legend. Error bars or shaded areas represent the standard error of the mean. *, **, and *** indicate p ≤ 0.05, p ≤ 0.01, and p ≤ 0.001, respectively.

Dithering strongly suppressed half-harmonic synchronisation to photic stimulation compared to periodic stimulation at *f*_max 1:2_, even for the intermediate dithering level (Fig 3C-D). This effect was manifest both for a global PLV measure (Fig 3C), where phase synchronisation is computed for 10 s trials, and a windowed PLV measure (Fig 3D), which is also sensitive to transient phase synchronisation (in both cases: p < 0.001 for both dithering levels, one-tailed). Moreover, synchronisation was reduced at the half-harmonic of stimulation more than at the stimulation frequency (comparison of ratios relative to the periodic condition with full modulation depth at 1:1 and 1:2): for the global PLV measure, p = 0.032 for *ζ* = 4.3%, and p = 0.005 for *ζ* = 9.2%, while for the windowed PLV, p = 0.001 for both dithering levels (one-tailed tests). The periodic condition with reduced modulation depth also showed a significant suppression of half-harmonic synchronisation, and a greater reduction in synchronisation at the half-harmonic of stimulation than at stimulation frequency as measured by both global and windowed PLVs (p ≤ 0.002 for all four tests, one-tailed). While the periodic condition with reduced modulation depth better preserved synchronisation at the stimulation frequency as measured by the global PLV (Fig 3C) than dithering, the intermediate dithering level preserved synchronisation at the stimulation frequency measured by the windowed PLV (Fig 3D) similarly to the periodic condition with reduced modulation depth (p = 0.38, two-tailed), and similarly suppressed 1:2 synchronisation (p = 0.74, two-tailed). The global PLV measure for the intermediate dithering level was also above that of the control condition (noise level) at the stimulation frequency, p < 0.001 (one-tailed), Fig 3C.

### 2.2 Half-harmonic responses are better explained by half-harmon entrainment than the superposition of evoked responses

To elucidate the nature of these half-harmonic responses, we generated synthetic data using averaged evoked potentials superimposed onto EEG control data according to the timing of stimulation in each trial and for each participant (Fig 4B). The superposition of evoked potentials hypothesis could account for the PLV values obtained in the data at the stimulation frequency (see 1:1 in Fig 4C1-2 and D1-2) as well as superharmonics for 10 and 20 Hz stimulation (see Fig S.3 in the supplementary material). However, regardless of the type of PLV triggers used (see Fig 4A), the linear superposition of evoked potentials could not account for the presence of half-harmonic synchronisation as measured by the PLV (compare Fig 4C2 and D2 to C1 and D1). Beyond linear superposition, evoked potentials could be modulated by non-linear sensory mechanisms such as saturation or gain control [32, 33]. To account for this, we additionally modulated evoked potentials at half the stimulation frequency. Such synthetic data could reproduce the response observed for the periodic condition with full modulation depth, but not for the dithered conditions (Fig 4C3 and D3). When using dithered PLV triggers, the discrepancy with the data was striking for both dithering levels (compare Fig 4D1 and D3). Similar results were obtained when using the averaged flash visual evoked potential (VEP) recorded in one participant (Fig S.4 in the supplementary material), or averaged evoked potentials including frequency components at half the stimulation frequency (Fig S.5 in the supplementary material).

**Figure 4:**
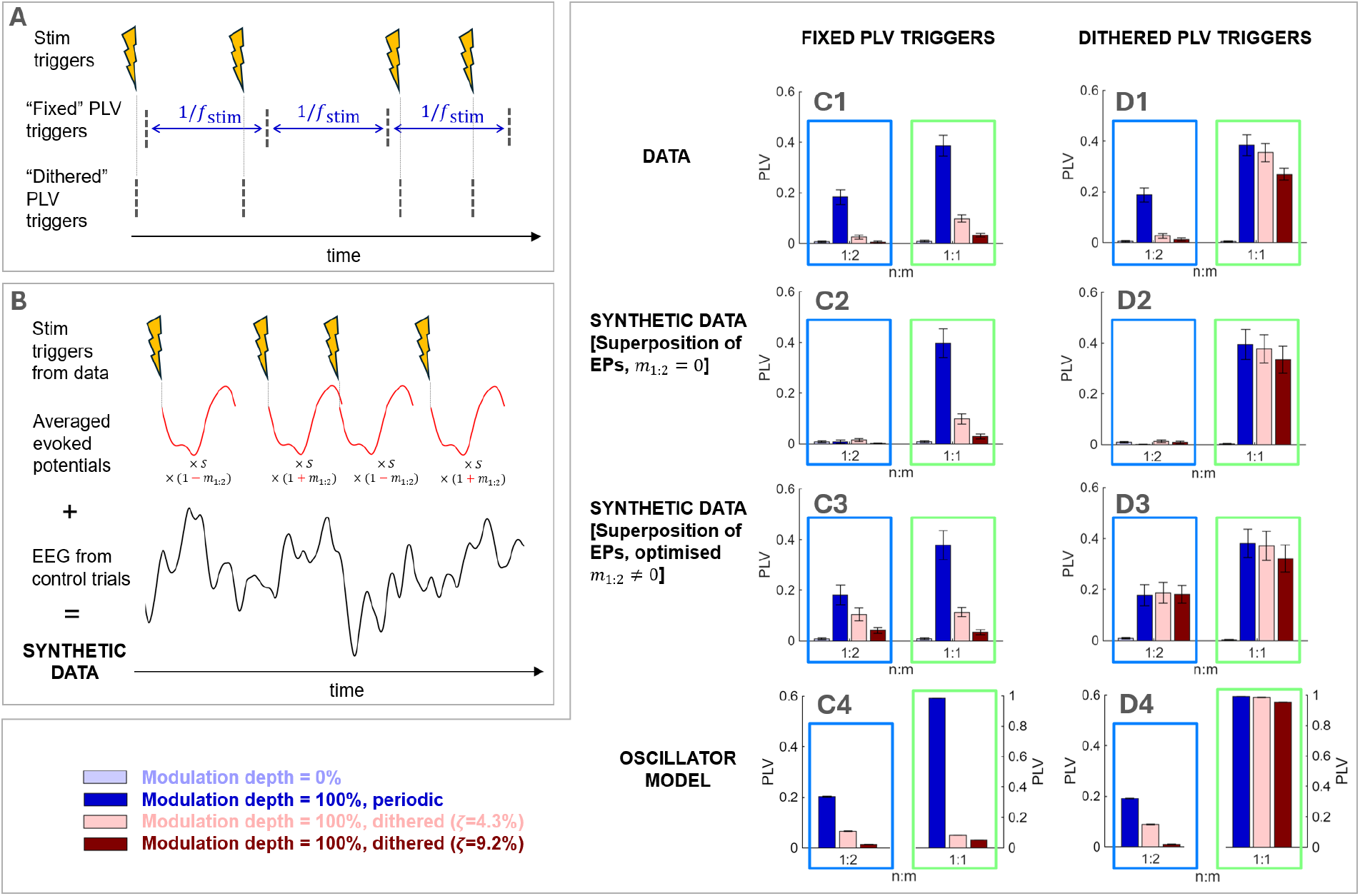
Half-harmonic responses to photic stimulation are better explained by half-harmonic entrainment than the superposition of evoked responses. **A**: Schematic of the difference between fixed PLV triggers and dithered PLV triggers (*f*_stim_ is the stimulation frequency). **B**: Schematic illustrating how synthetic data is generated according to the superposition of evoked potentials hypothesis. For each participant, the scale factor *S* was determined to match the data PLV_1:1_ for the periodic condition. In C3 and D3, the modulation factor *m*_1:2_ was chosen to match the data PLV_1:2_ for the periodic condition for each participant. **C-D**: PLV at the stimulation frequency (1:1) and its half-harmonic (1:2) at the group level, for fixed PLV triggers (C) and dithered PLV triggers (D). Comparison between the data, synthetic data generated according to the superposition of evoked potentials hypothesis (without and with modulation at the half-frequency), and an oscillator model (sine circle map with noise). Panels C-D share the same legend, and error bars represent the standard error of the mean (error bars are hardly visible in C4 and D4 due to low cross-trial variability).

We investigated the alternative hypothesis of subharmonic entrainment using an oscillator model. We simulated the sine circle map, which is the simplest model describing the influence of periodic stimulation on a single neural oscillator, with the addition of noise to the oscillator’s frequency to approximately match 1:2 synchronisation for periodic stimulation to the data. The resulting PLV_1:2_ for dithered stimulation are more consistent with the data for both types of PLV triggers (Fig 4C4-D4) than the PLV_1:2_ based on the superposition hypothesis, in particular for dithered PLV triggers.

## 3 Discussion

In this study, we showed using photic stimulation and EEG recordings in healthy participants that slightly jittering stimulation pulses (dithered stimulation) suppresses half-harmonic synchronisation (Fig 3) relative to perfectly periodic flicker at the individual half-harmonic optimum frequency (*f*_max 1:2_), as previously predicted by mathematical modelling [19]. Notably, dithered stimulation suppressed half-harmonic synchronisation more than synchronisation at the stimulation frequency (in particular when using a windowed measure of synchronisation, Fig 3D). This was also the case for the periodic condition with reduced stimulation depth, as predicted by theory.

Additionally, we demonstrated by generating synthetic data that the half-harmonic responses observed in participants cannot be explained by the superposition of evoked responses, even when evoked responses are modulated at the half-harmonic frequency (Fig 4C2-3, D2-3). The discrepancy with data was the largest when the PLV was computed with dithered PLV triggers that follow dithered stimulation timing variability. Instead, the half-harmonic responses of a minimal oscillator model receiving periodic and dithered stimulation better matched the group level data (Fig 4C4, D4), suggesting the presence of half-harmonic entrainment.

Together, these findings are consistent with the view that half-harmonic EEG synchronisation during photic stimulation predominantly reflects half-harmonic entrainment rather than the summation of evoked responses, and that dithering, as well as reducing stimulation amplitude, effectively suppress subharmonic entrainment.

### 3.1 Limitations

The sample size is limited (n=10), in part due to the rejection of datasets with insufficient half-harmonic response to periodic stimulation. Nonetheless, the measured effect size is large (Cohen’s d for the difference between PLV_1:2_ for periodic stimulation with full modulation depth and the first/second dithering level is 1.77/2.06, respectively). While dithering also significantly suppressed superharmonic synchronisation, and more so than synchronisation at the stimulation frequency (Fig S.3B1, C1 in the supplementary material, p-values in the caption), we could not conclude on the potential of dithering to suppress superharmonic entrainment because we could not distinguish these superharmonic responses from the superposition of evoked potentials (Fig S.3). Due to experimental time constraints, we were also unable to test stimulation conditions that may better preserve the 1:1 response, in particular lower dithering levels than *ζ* = 4.3%, and the combination of low dithering levels with reduced modulation depth. Such stimulation conditions may be as good or better than the periodic condition with 66% modulation depth at suppressing 1:2 while preserving 1:1 synchronisation. Finally, to assess whether the half-harmonic data may be more consistent with entrainment, we intentionally simulated a minimal (phase-only) oscillator model. More realistic models (e.g. coupled oscillators or neural mass models [13, 29, 27]) may better reproduce the data.

### 3.2 Half-harmonic responses are consistent with entrainment, not evoked-potential superposition

The nature of the 1:1 response to rhythmic sensory stimulation such as photic stimulation is still debated, with some studies pointing to a simple superposition of evoked responses [20, 34], and others to neural entrainment and resonance of neural circuits [21, 24, 22, 23]. Half-harmonic responses have been reported in humans [25, 26, 27, 28, 6, 29] and animals [32, 35] in response to photic stimulation, however their underlying mechanism has received little attention. While we could not distinguish in our data 1:1 and superharmonic responses from a superposition of evoked responses, we provided evidence that half-harmonic responses are inconsistent with the superposition of evoked responses hypothesis and may represent neural entrainment. Patient-specific synthetic data generated according to the superposition of evoked responses hypothesis failed to account for any 1:2 synchronisation. When adding an explicit half-frequency modulation of evoked responses (which could be the result of non-linear sensory mechanisms), the synthetic data could match the PLV_1:2_ in response to periodic stimulation yet could not reproduce the PLV_1:2_ observed for dithered stimulation, in particular for dithered PLV triggers (Fig 4). We also note that since the illumination and modulation depth were the same for the periodic and dithered stimulation used in this analysis and the pulse timing differences under the dithered conditions are minimal, it is unlikely that a potential non-linear sensory mechanism would behave differently under the periodic and dithered conditions. Moreover, supplementary analyses that 1) substituted the averaged evoked potential for a flash VEP measured separately or 2) constructed averaged responses that already contain half-frequency components from the data led to the same mismatch under dithering.

Conversely, the oscillator model reproduced the drop in PLV_1:2_ as *ζ* increases for dithered PLV triggers. Similarly to the data, the oscillator’s half-harmonic responses to dithered stimulation were not tightly temporally locked to every other stimulation trigger as revealed by PLV_1:2_ with dithered PLV triggers (dithering had no impact on PLV_1:2_ in the superposition hypothesis even with *m*_1:2_ ≠ 0, Fig 4D3). While the presence of an on-going oscillation is sometimes cited as a prerequisite for entrainment [36, 37], recent data shows that sub-harmonic entrainment does not necessarily require an on-going oscillation that can be measured in the absence of stimulation, even when using invasive recordings [17]. Such a pre-existing oscillation could facilitate entrainment [38], however unsynchronised oscillators with the capacity to synchronise may be sufficient.

Together, the evidence we present weigh against the superposition hypothesis for half-harmonic responses and favor a half-harmonic entrainment mechanism sensitive to the temporal statistics of the pulse train beyond simple locked responses.

### 3.3 A simple design rule to quench harmonic synchronisation: add jitter

We developed dithered stimulation as a method to suppress sub- and superharmonic entrainment using mathematical modelling [19]. Here, we confirmed that adding a modest amount of jitter to the timing of stimulation pulses (*ζ* = 4.3%) can effectively suppress subharmonic synchronisation in response to sensory stimulation in humans, while maintaining some level of 1:1 synchronisation. Another study provided evidence that poisson stimulation trains prevent superharmonic responses during optogenetic excitatory stimulation in mice [39], with a broad low power 1:1 response likely due to the large pulse timing variability. It was recently suggested that half-harmonic entrainment to brain stimulation can functionally disconnect neural oscillations [17, 15]. When such disconnection could lead to deleterious effects, dithering offers a practical engineering principle to avoid half-harmonic synchronisation.

Given that dithering or other types of noisy pulse trains have been reported to suppress harmonic responses in minimal and more realistic computational models [19], in mice using optonogenetics [39], and in humans using photic stimulation (this study), the efficacy of dithering in suppressing harmonic responses may generalise to other stimulation modalities. Such modalities may include transcranial alternating current stimulation, which is known to entrain neural oscillations [40], and DBS, which was recently shown to entrain neural oscillations at the half-harmonic of the stimulation frequency [11, 12, 13]. This observation has gathered a significant amount of interest, in part because of the potential involvement of half-harmonic entrainment in the therapeutic effects of DBS in movement disorders [14, 15, 16]. Since dithered stimulation can modulate half-harmonic synchronisation, it could provide a means to causally investigate the therapeutic relevance of half-harmonic synchronisation. It is however unclear how patient groups may react to dithering compared to healthy participants, and this should be the subject of future work. From a practical perspective, dithered stimulation can be implemented in neurostimulators with limited capabilities by toggling within a finite set of stimulation frequencies [19]

### 3.4 Dithering vs reducing stimulation amplitude

Theory also predicts that sufficiently reducing stimulation amplitude (which corresponds to modulation depth in this study) will suppress 1:2 entrainment, and this was confirmed by our results. When considering the global PLV, the periodic condition with reduced modulation depth suppressed half-harmonic synchronisation similarly to the intermediate dithering level, but better preserved synchronisation at the stimulation frequency. However, results were comparable when considering the windowed PLV. Reducing stimulation amplitude impacts 1:2 entrainment through a different mechanism than dithering: the system simply leaves the 1:2 Arnold tongue (see Fig 1B1). The efficacy of a reduced stimulation amplitude in suppressing 1:2 synchronisation strongly depends on the system of interest (which determines the shape of the Arnold tongues) and stimulation parameters (where the system is with respect to the boundaries of the 1:2 and 1:1 Arnold tongues).

Whether reducing stimulation amplitude is a better choice than dithering to suppress 1:2 synchronisation while preserving 1:1 synchronisation will therefore depend on the system of interest. In some cases, it may be optimal to combine dithering with a reduced stimulation amplitude. Reducing stimulation amplitude may not always be possible, for example when supra-threshold stimulation is needed, when a certain amount of energy needs to be provided for therapeutic effect, or due to neurostimulator limitations. In such cases, manipulating the timing of stimulation pulses, as with dithering, would be the only way to suppress harmonic synchronisation.

### 3.5 Conclusion

In summary, we demonstrated that dithering can suppress half-harmonic synchronisation to photic stimulation in humans. Moreover, we provided evidence that these half-harmonic responses are consistent with entrainment but not with the evoked potential superposition hypothesis. When stimulation amplitude cannot be reduced, these findings support dithering as a simple open-loop design principle for selective entrainment, and highlight its potential to enable brain stimulation therapies where there are physiological rhythms to reinforce and pathological rhythms that should not be entrained. Dithering could also enable new mechanistic investigations through its ability to modulate 1:2 synchronisation while ensuring the delivered energy remains constant.

## 4 Methods

### 4.1 Participants

We recruited right-handed healthy participants, who were screened to minimise the risk of photic stimulation causing epileptic seizures. We included participants above 20 years old (to reduce risk of undiagnosed epilepsy) and below 60 years old, see Table A in the supplementary material for detailed inclusion/exclusion criteria. The study was approved by the Medical Sciences Interdivisional Research Ethics Committee of the University of Oxford (approval reference: R93268/RE001), and all participants gave written informed consent. We recorded data from 16 participants (8 female, mean age = 31.6 ± 10.0 years old, see Table B in the supplementary material).

### 4.2 Photic stimulation and EEG setup

Photic stimulation was delivered via a white light emitting diode (LED) strip with a diffuser in front. The LED strip was driven by a Teensy 4.1 microcontroller, allowing precise control of the stimulation waveform. The microcontroller’s voltage output was converted into a current waveform using a custom control board (Fig S.1A in the supplementary material). Participants were sat approximately 75 cm away from the photic stimulation device with the device at eye level. The illuminance at the level of the participants’ eyes was approximately 70 lux (during continuous illumination, measured using an RS-92 light meter, RS PRO). The light flicker was presented as square pulses (20% duty cycle) since square pulses evoke a stronger response at the stimulation frequency than sinusoidal waveforms [41], and were also found in pilot testing to elicit a stronger subharmonic response. A photodiode placed on the LED panel allowed synchronised acquisition of the illumination waveform and EEG signal. The non-linearity of the photodiode was corrected using a calibration procedure (see Fig S.2 in the supplementary material for more details).

We recorded EEG using a TMSi (Twente Medical Systems) amplifier and a 10-10 EEG cap with 64 electrodes. Only 15 electrodes were used (see Fig S.1B in the supplementary material), located more posteriorly towards the visual cortex and referenced using common average referencing. The AFz electrode was used as a ground electrode. Impedances were kept under 5 kW. EEG data were recorded with a sampling rate of 4096 Hz.

### 4.3 Experimental paradigm

To identify the stimulation frequency leading to the largest power response at the half-harmonic (denoted *f*_max 1:2_), participants were first exposed to a frequency sweep with periodic flicker (*ζ* = 0%) from 15 Hz to 43 Hz in 2 Hz increments (Fig 2A1), with two repeats per frequency. This range was previously found to elicit half-harmonic responses in most subjects [28]. Modulation depth, defined as (*I*_max_ − *I*_min_) */I*_max_ where *I*_max_ is the illuminance during the “on” part of the pulse, and *I*_min_ is the illuminance during the “off” part of the pulse (see Fig 2C)), was 100%. The power of the EEG response was obtained for each stimulation frequency using Welch’s method (8 segments per trial with 50% overlap), and averaged across trials and EEG channels. The stimulation frequency with the largest averaged power response at the half-harmonic of the stimulation frequency was identified as *f*_max 1:2_ (see example in Fig S.1D in the supplementary material).

Next, periodic stimulation was compared to dithered stimulation by exposing participants to flicker of frequency *f*_max 1:2_, but also 10, and 20 Hz. For each stimulation frequency, periodic conditions (*ζ* = 0%) with modulation depths of 0% (control condition with no flicker), 66%, and 100% were included, as well as two dithered conditions (*ζ* = 4.3% and *ζ* = 9.2%, with a modulation depth of 100% in each case) – see Fig 2A2. Each stimulation condition was repeated four times. The manuscript focuses on stimula-tion at *f*_max 1:2_, but results for 10 Hz and 20 Hz stimulation are included in the supplementary material.

During both experimental parts, the order in which stimulation conditions (including repeats) were presented was randomised. Stimulation periods lasted 10 s and were followed by 20 s rest period (25 s during the frequency sweep).

### 4.4 Quantifying synchronisation

A coarse assessment of synchronisation was first performed by computing the power of the EEG response. All trials were visually inspected for artefacts and trials with artefacts such as motion, muscle, or electrode pop artefacts were not included in the analysis. We trimmed 200 ms off the beginning and the end of each trial to remove potential onset and offset effects. The power of the EEG response was obtained for each stimulation frequency using Welch’s method (8 segments per trial with 50% overlap), and averaged across trials and EEG channels.

To better quantify the degree of EEG signal synchronisation to stimulation, we computed the phase-locking value (PLV), which measures the concentration of the EEG signal’s phase according to the timing of stimulation. We considered the PLV at integer ratios of the stimulation frequency (1:1 and 1:2 in the main text, as well as 2:1 and 3:1 in the supplementary material). For each n:m ratio considered, the EEG data was bandpass filtered around *f*_filt_ = (*n/m*) *f*_stim_, with a half-width relative to the filter’s center frequency given by *df* = (*n/m*) *f*_stim_*/*10, corresponding to *df* = 1 Hz for *f*_stim_ = 10 Hz at 1:1. We applied a second order butterworth filter both forwards and backwards to minimise phase distortion, and obtained the Hilbert phase *ψ*_n:m_ for each n:m ratio considered. For each trial, and each EEG channel, the PLV was computed for n:1 ratios as

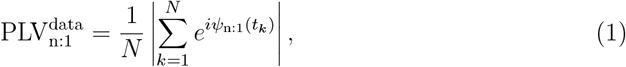

where *N* is the number of pulses in the trial considered, *ψ*_n:1_(*t*) is the Hilbert phase obtained from the corresponding filtered EEG signal, and the *t*_*k*_’s are the times of the PLV triggers, which are described below. The PLV was also computed for 1:2 by considering only every other trigger using

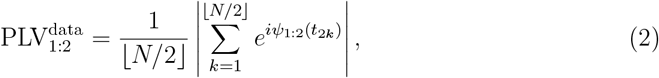

with. the floor function. PLV values were averaged over channels and non-rejected trials, with averaged values denoted 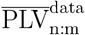.

We computed the PLV using two types of PLV triggers, which are the same for periodic stimulation but not for dithered stimulation (Fig 4A). The first type, which we call “fixed PLV triggers”, was used to assess synchronisation at the average stimulation frequency, and employed as PLV triggers the times of stimulation in the absence of dithering. For *ζ* = 0%, we used the times of stimulation from the current trial, whereas for *ζ >* 0%, we used the times of stimulation from the first non-rejected trial with the same stimulation frequency and modulation depth but with *ζ* = 0%. The second type, which we call “dithered PLV triggers”, was used to contrast the data with various models, and employed as PLV triggers the times of stimulation in the current trial regardless of *ζ*. In both cases, the times of stimulation were identified through threshold crossings of pulses’ leading edges in the illumination signal obtained from the photodiode.

In all cases, to remove from PLV estimates the contribution of spurious phase-locking due to phase alignment with stimulation happening by chance, we computed the PLV for pink noise signals put through the exact same analysis as the data, including filtering (using the same PLV triggers used for the data). The PLV estimate with the noise contribution removed is given by

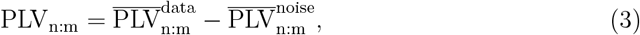

where 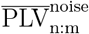 is PLV for noise computed as for the data (using equations (1) or (2)), averaged as for the data.

To increase sensitivity to transient locking which may be present in the data, we also computed a windowed version of the PLV, whereby we obtained the PLV in windows of duration relative to the filter’s center frequency given by 500 × 40*/f*_filt_ ms (500 ms for *f*_filt_ = 40 Hz), with overlap also relative to the filter’s center frequency given by 200 × 40*/f*_filt_ ms (200 ms for *f*_filt_ = 40 Hz), and averaged the resulting values. We denote the windowed PLV by 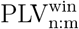 (with noise removed as per equation (3)).

We excluded from the analysis five participants who had a very weak response at the half-harmonic of stimulation 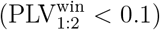 in the periodic condition for *f*_stim_ = *f*_max 1:2_ and a modulation depth of 100%. We also excluded from the analysis one participant who did not have at least two non-rejected trials for each stimulation condition for *f*_stim_ = *f*_max 1:2_.

### 4.5 Synthetic data based on the superposition hypothesis

To assess whether the PLVs observed in data at the 1:2 subharmonic could be accounted for simply by sensory evoked potentials, synthetic data was generated and put through the same analysis pipeline as the data. For each participant included in the analysis, we generated synthetic data by 1) replacing each stimulation trial by EEG data corresponding to one of the participant’s randomly selected control trials (where the LED panel was on but not flashing), and 2) aligned to each stimulation trigger recorded from the photodiode in the stimulation trial, summing scaled averaged evoked potentials to the control EEG data (Fig 4B). Averaged evoked potential were obtained by averaging all 1*/f*_stim_ long epochs directly following stimulation triggers in trials at the stimulation frequency considered, with *ζ* = 0%, and a modulation depth of 100%. This was done independently for each EEG channel and each subject, using the corresponding EEG data high-passed at 1Hz, and low-passed at 80Hz, with a notch filter at 50Hz (line noise in the UK). Other types of averaged evoked potentials were also considered as described in the supplementary methods. The scale factor *S* was chosen to match the channel-average PLV at the stimulation frequency, *ζ* = 0%, and 100% modulation depth in each participant’s data and synthetic data.

We also considered the possibility that sensory evoked potentials could be modulated at half the stimulation frequency by non-linear sensory mechanisms such as saturation or gain control [32, 33]. For example, the gain of the response could be reduced for a short period after a strong flash. To account for such a potential mechanism, we alternated between modulating consecutive evoked potentials by the factor 1 + *m*_1:2_, and the factor 1−*m*_1:2_ (Fig 4B). We chose *m*_1:2_ to match the channel-average PLV at half the stimulation frequency, *ζ* = 0%, and 100% modulation depth in each participant’s data and synthetic data.

### 4.6 Oscillator model

To investigate whether the PLV levels observed in the data at the 1:2 subharmonic may be better accounted for by entrainment of an oscillator, we simulated a simple oscillator model. As in [19], we considered the sine circle map, which is the simplest model describing the influence of periodic stimulation on a single neural oscillator. Entrainment can arise because a stimulus may advance or delay the phase of an oscillator depending on the phase at which it is applied. This concept is captured by the phase response curve (PRC) of the oscillator, which describes the change in phase of the oscillator as a function of the stimulation phase. The PRC of the sine circle map is a simple sinusoid (*Z*(*θ*) = sin *θ*). The model we used maps the phase of an oscillator right before stimulation pulse *n* (denoted *θ*_*n*_) to the phase of the oscillator right before stimulation pulse *n* + 1 (denoted *θ*_*n*+1_). The map is described by

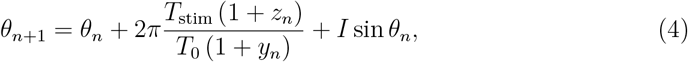

where *I* is the stimulation magnitude and, to model dithered stimulation, the stimulation period *T*_stim_ = 1*/f*_stim_ is multiplied by (1 + *z*_*n*_), with *z*_*n*_ normal random numbers sampled from 𝒩 (0, *ζ*^2^), and *ζ* the dithering level. We have also added noise to the oscillator’s natural period *T*_0_ to allow us to reduce the PLV at the 1:2 subharmonic in the absence of dithering to the average value observed in the data. The oscillator’s noise level is denoted *ζ*_mod_, and *y*_*n*_ are normal random numbers sampled from 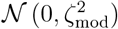.

For comparison with the data group average results, we picked *f*_stim_ = 30 Hz (close to the group average of *f*_max 1:2_). We matched the PLV at the 1:2 subharmonic in the absence of dithering to the average value observed in the data by using *ζ*_mod_ = 0.083. For each dithering level used in the data, we simulated 40 trials of 6000 pulses each. To calculate the PLV using fixed PLV triggers, we sampled the phase at *dt* = 0.0001 s between points given by the map (assuming the oscillator’s frequency stays constant between stimulation pulses). Because the PLV_1:2_ metric is also sensitive to PLV at the 1:1 ratio in the absence of filtering (which, contrary to the data, cannot be applied here since the oscillator only has a phase and no amplitude), a PLV_1:2_ metric comparable to the data was obtained by subtracting PLV_1:1_ to PLV_1:2_ (both computed at the same oscillator natural frequency and amplitude). This process also removes spurious synchronisation due to chance. We averaged these results within a region of interest focused on the center of the 1:2 tongue (natural frequency 15 Hz, and stimulation magnitude between 0.85 and 1.35 a.u.).

### 4.7 Statistical tests

To compare synchronisation between conditions at the group level, we performed signed rank tests for paired data (n = 10 pairs for all tests). All statistical tests in this study were performed under false discovery rate (FDR) control at 5% according to the Benjamini and Hochberg procedure [42]. All p-values < 0.05 were found to also be significant under FDR control.

## Funding information

R.B. was supported by Medical Research Council grant MC_UU_00003/1. B.D. was jointly supported by the Royal Academy of Engineering and Rosetrees under the Research Fellowship programme. S.H., A.P., and H.T. were supported by the Medical Research Council (MRC) (MC_UU_00003/2), the Medical and Life Sciences Translational Fund (MLSTF) from the University of Oxford, the National Institute for Health and Care Research (NIHR) Oxford Biomedical Research Centre, and the Rosetrees Trust, UK. S.H. was also supported by a Non-Clinical Postdoctoral Fellowship from the Guarantors of Brain, an International Exchanges Award (IES\R3\213,123) from The Royal Society, and a Senior Research Fellowship from Parkinson’s UK. A.S. and N.S. were supported by the Medical Research Council grant MC_UU_00003/6. T.D. was supported by the Royal Academy of Engineering and the NIHR Invention for Innovation Programme.

## Data availability statement

The data collected as part of this work will be shared openly before publication.

## Notes

### Competing Interest Statement

B.D., T.D., and R.B. are stakeholders in an intellectual property application based on this work.

