## Supplementary material for "Dithering suppresses half-harmonic neural synchronisation to photic stimulation in humans"

### A Supplementary methods: additional synthetic data with superposition of evoked potentials

To assess whether the PLVs observed in data at the 1:2 subharmonic could be accounted for simply by sensory evoked potentials, we also generated synthetic data using two other types of averaged evoked potentials.

In figure S.4, we used the averaged flash VEP obtained by averaging the EEG response to 45 flashes (two-second interval between flashes) at 100% modulation depth in one participant (participant 10). This was done independently for each EEG channel, and the EEG was band-pass filtered between 1 and 45 Hz before averaging. This averaged flash VEP was truncated at 40 0ms, and was used for all participants. The scale factor $S$, and when applicable the modulation factor $m_{1:2}$, were determined independently for each subject as before.

In figure S.5, we used averaged evoked potentials including frequency components at half the stimulation frequency. These averaged evoked potentials were obtained by averaging $2/f_{\text{stim}}$ long epochs directly following every other stimulation triggers in trials at the stimulation frequency considered, with $\zeta = 0\%$, and a modulation depth of 100%. This was done independently for each EEG channel and each participant, using the corresponding EEG data high-passed at 1 Hz, and low-passed at 80 Hz, with a notch filter at 50 Hz. The scale factor $S$, and when applicable the modulation factor $m_{1:2}$, were determined independently for each subject as before. In the perfectly periodic case and with $m_{1:2} = 0$, frequency components in the averaged evoked potential at half the stimulation frequency cancel out. However this is in general not the case with dithering, and this experiment allowed us to investigate whether interactions between $m_{1:2} \neq 0$ and frequency components in the averaged evoked potential at half the stimulation frequency might give rise to the PLV pattern observed in the data with dithered stimulation.

### B Supplementary tables

| | |
|---|---|
| Inclusion criteria | • Participant is willing and able to give informed consent for participation in the study<br>• Participant is mobile and can come to the lab without assistance<br>• Right-handed<br>• 20 to 60 years old<br>• Fluent in English<br>• Normal or corrected to normal vision<br>• No current significant medical condition<br>• No personal and/or family history of epilepsy<br>• Not currently taking any medications (except the contraceptive pill)<br>• Not pregnant or planning to get pregnant for the duration of the study |
| Exclusion criteria | • History of neurological conditions or currently diagnosed with a neurological condition (including paralysis, muscle weakness, poor coordination, loss of sensation, seizures, confusion, pain, and altered levels of consciousness)<br>• Any kind of epilepsy (including photosensitive epilepsy), or a family history of any kind of epilepsy.<br>• Any history or family history of seizures (including febrile convulsions in childhood)<br>• Any known significant sensitivity to flashing lights, or previous adverse experience in response to flashes, alternating patterns, or stripes of contrasting colours (for example from television or computer screen, strobe lights, flickering natural light such as sunlight flickering through trees...)<br>• History of migraines<br>• History of psychiatric conditions or currently diagnosed with a psychiatric condition, including bipolar disorder, depression or anxiety disorders<br>• Current consumption of any medication (except the contraceptive pill), including psychotropic drugs (such as antidepressants and neuroleptics)<br>• Eye disease (including glaucoma, retinitis, retinopathy, or macular degeneration)<br>• Any cardiac pathology<br>• Planning to get pregnant or likely to be pregnant during the course of the study<br>• Participant slept at least one hour less than their normal sleeping duration before the experimental session.<br>• Consumption of 3 or more units of alcohol in the last 24 hours<br>• Withdrawal from regular alcohol consumption<br>• High consumption of caffeine (more than one cup of coffee, or other source of caffeine in the last hour)<br>• Consumption of recreational drugs in the last 24 hours |

**Table A: Inclusion and exclusion criteria used in the study.**

| participant ID | gender | age | $f_{\text{max } 1:2}$ (Hz) | $\text{PLV}^{\text{win}}_{1:2}$ | included in analysis |
|---|---|---|---|---|---|
| 1 | F | 21 | 31 | 0.255 | yes |
| 2 | F | 21 | 43 | 0.081 | no |
| 3 | M | 52 | 31 | 0.145 | yes |
| 4 | F | 35 | 43 | 0.021 | no |
| 5 | M | 29 | 43 | 0.216 | yes |
| 6 | F | 22 | 37 | 0.026 | no |
| 7 | M | 51 | 39 | NA | no |
| 8 | M | 41 | 39 | 0.024 | no |
| 9 | F | 20 | 39 | 0.113 | yes |
| 10 | M | 38 | 39 | 0.137 | yes |
| 11 | M | 31 | 19 | 0.102 | yes |
| 12 | F | 28 | 33 | 0.133 | yes |
| 13 | M | 28 | 29 | 0.184 | yes |
| 14 | M | 35 | 31 | 0.182 | yes |
| 15 | F | 23 | 33 | 0.036 | no |
| 16 | F | 31 | 29 | 0.182 | yes |

**Table B: Participant demographics and inclusion for analysis based on EEG response.** Out of 16 recorded datasets, 10 were included in the study. The fourth column gives the stimulation frequency identified during the initial frequency sweep as giving rise to the largest power response at the half-harmonic frequency ($f_{\text{max } 1:2}$). The fifth column gives $\text{PLV}^{\text{win}}_{1:2}$ for $f_{\text{stim}} = f_{\text{max } 1:2}$, $\zeta = 0\%$, and a modulation depth of 100%. The $\text{PLV}^{\text{win}}_{1:2}$ entry for participant 7 reads NA (not available) because all trials were rejected for $f_{\text{stim}} = f_{\text{max } 1:2}$, $\zeta = 0\%$, and a modulation depth of 100%.

### C Supplementary figures



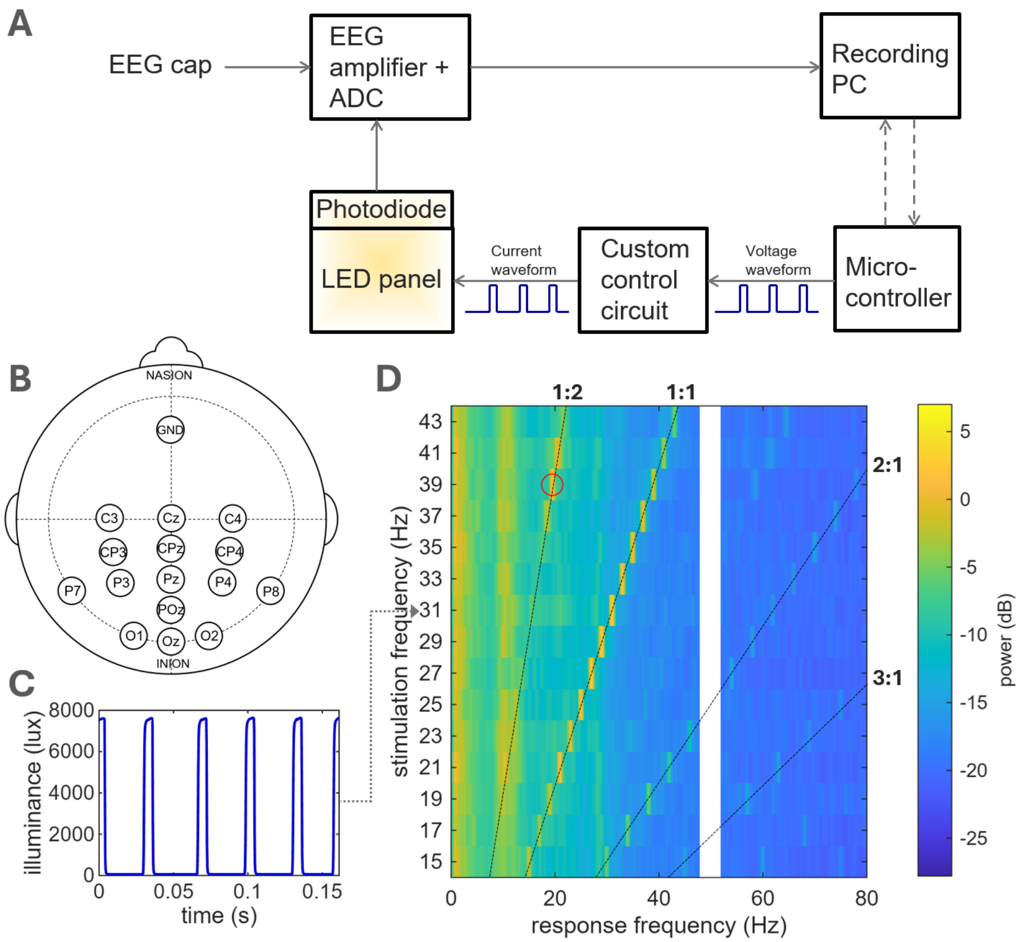


**Figure S.1: Measuring EEG responses to photic stimulation.** **A**: Sketch of the experimental setup. Dashed arrows indicate intermittent communication between devices. **B**: EEG electrodes used in the study. **C**: Example photodiode output during 31 Hz periodic stimulation (corrected using the calibration presented in Fig S.2). **D**: EEG responses to periodic stimulation (full modulation depth) averaged over channels for stimulation frequencies between 15 and 43 Hz in one participant. The colorbar shows EEG power averaged over channels, such that each row of this plot represents the EEG power spectrum for the corresponding stimulation frequency on the vertical axis. The response at 50Hz is hidden due to line noise. The alpha rhythm is visible around a response frequency of 10 Hz. Dashed black lines highlight responses at integer ratios of the stimulation frequency. The maximum response at half the stimulation frequency is obtained for a stimulation frequency of $f_{\text{max } 1:2} = 39$ Hz and is indicated by a red circle.

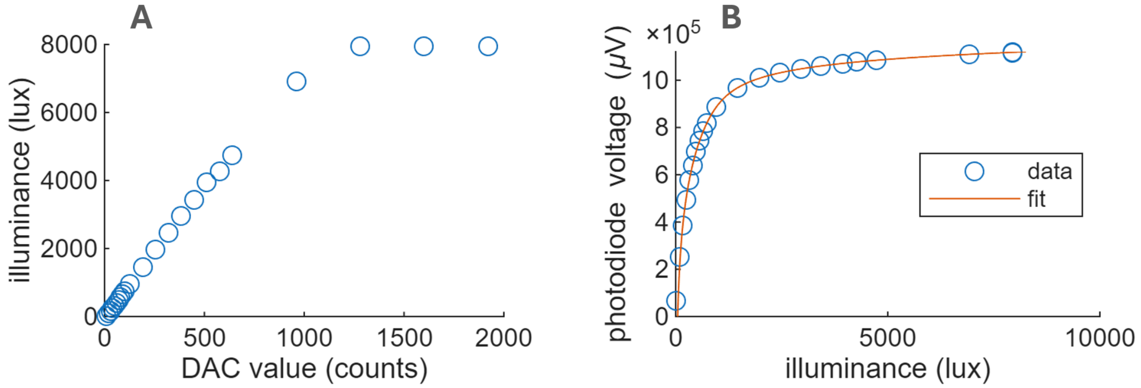


**Figure S.2: LED panel and photodiode calibration.** **A**: Illuminance measured at the LED panel's diffuser as a function of the digital-to-analog converter (DAC) value set in the microcontroller. This curve characterises the LED panel's response. The maximum DAC value used in this study is 1100 counts (just at the onset of saturation). **B**: Voltage measured by the photodiode at the LED panel's diffuser as a function of illuminance, also measured at the LED panel's diffuser. This curve characterises the photodiode's response. Data points are shown as blue circles, and the bi-exponential fit used to correct the output of the photodiode is shown in red. Illuminance was measured using a light meter (RS-92 Light Meter, RS PRO).

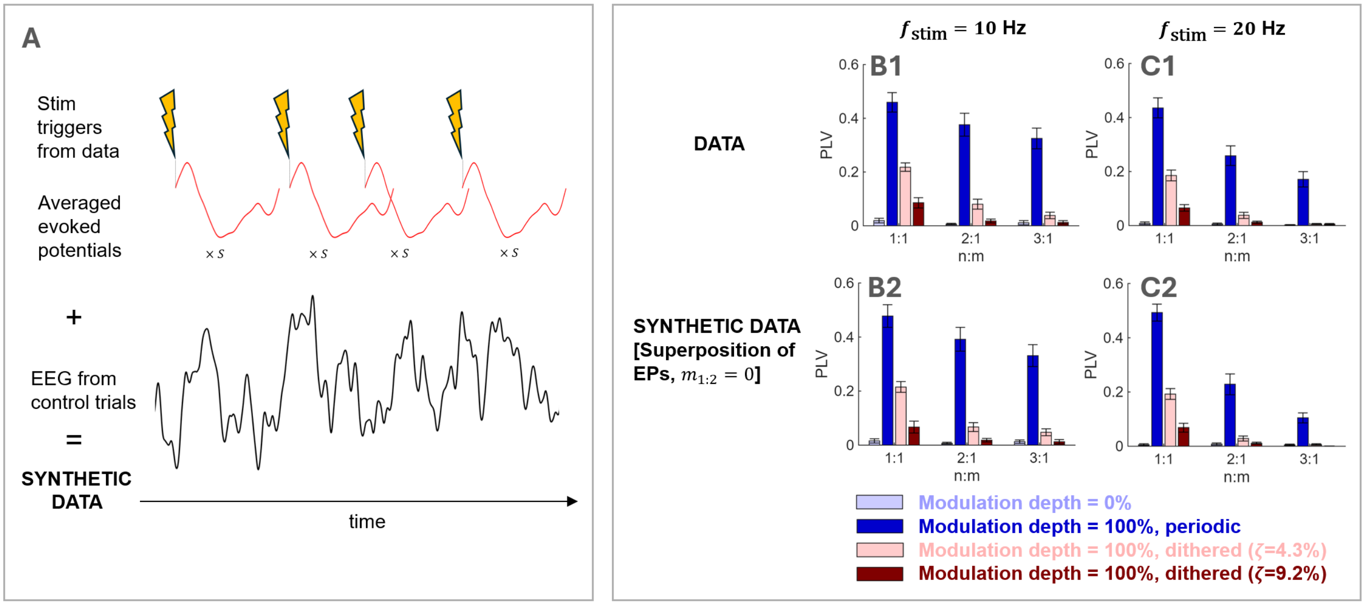


**Figure S.3: Superharmonic responses to photic stimulation can be reproduced by the superposition of evoked responses. A**: Schematic illustrating how synthetic data is generated according to the superposition of evoked potentials hypothesis. For each participant, the scale factor $S$ was determined to match the data $\text{PLV}_{1:1}$ for the periodic condition. **B-C**: PLV at the stimulation frequency (1:1) and two superharmonic (2:1 and 3:1) at the group level, using fixed PLV triggers. Comparison between the data and synthetic data generated according to the superposition of evoked potentials hypothesis. We note that superharmonic responses are strongly reduced by dithering in both cases. In the data (B1-C1), synchronisation was reduced at the superharmonics of stimulation more than at the stimulation frequency (comparison of ratios relative to the periodic condition with full modulation depth at 1:1 vs 2:1 and 3:1) For 10 Hz stimulation (B1): for 2:1 vs 1:1, $p = 0.001$ ($\zeta = 4.3\%$), and $p = 0.0039$ ($\zeta = 9.2\%$); for 3:1 vs 1:1, $p = 0.001$ ($\zeta = 4.3\%$), and $p = 0.0039$ ($\zeta = 9.2\%$). For 20 Hz stimulation (C1): for 2:1 vs 1:1, $p = 0.001$ ($\zeta = 4.3\%$), and $p = 0.002$ ($\zeta = 9.2\%$); for 3:1 vs 1:1, $p = 0.002$ ($\zeta = 4.3\%$), and $p = 0.0137$ ($\zeta = 9.2\%$). All these tests were one-tailed. Panels B-C share the same legend, and error bars represent the standard error of the mean. Note that these stimulation frequencies did not produce subharmonic responses to periodic stimulation with full modulation depth.

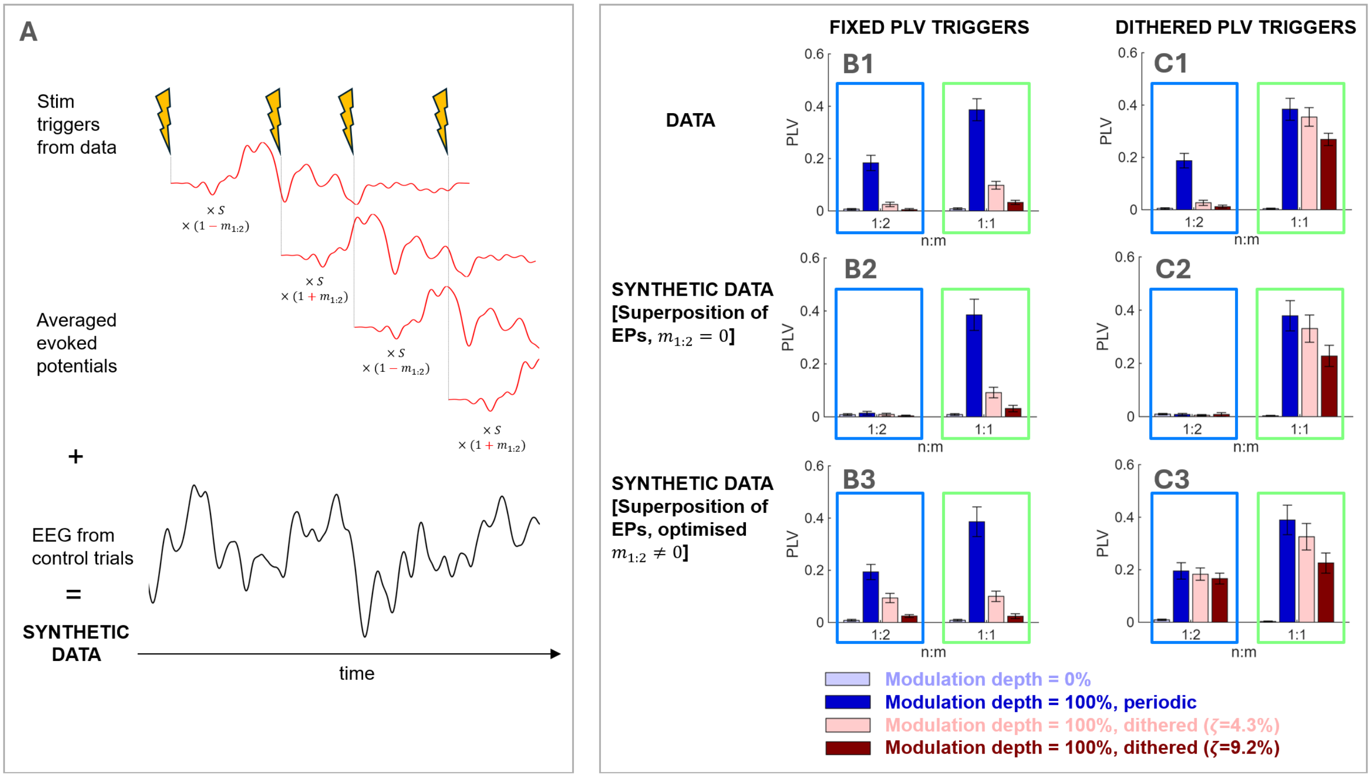


**Figure S.4: Half-harmonic responses to photic stimulation are inconsistent with the superposition of flash VEPs.** **A**: Schematic illustrating how synthetic data is generated according to the superposition of evoked potentials hypothesis, using the averaged flash VEP obtained in one participant. For each participant, the scale factor $S$ was determined to match the data $\text{PLV}_{1:1}$ for the periodic condition. In B3 and C3, the modulation factor $m_{1:2}$ was chosen to match the data $\text{PLV}_{1:2}$ for the periodic condition for each participant. **B-C**: PLV at the stimulation frequency (1:1) and its half-harmonic (1:2) at the group level, for fixed PLV triggers (B) and dithered PLV triggers (C). Comparison between the data and synthetic data generated according to the superposition of evoked potentials hypothesis (without and with modulation at the half-frequency). Panels B-C share the same legend, and error bars represent the standard error of the mean.

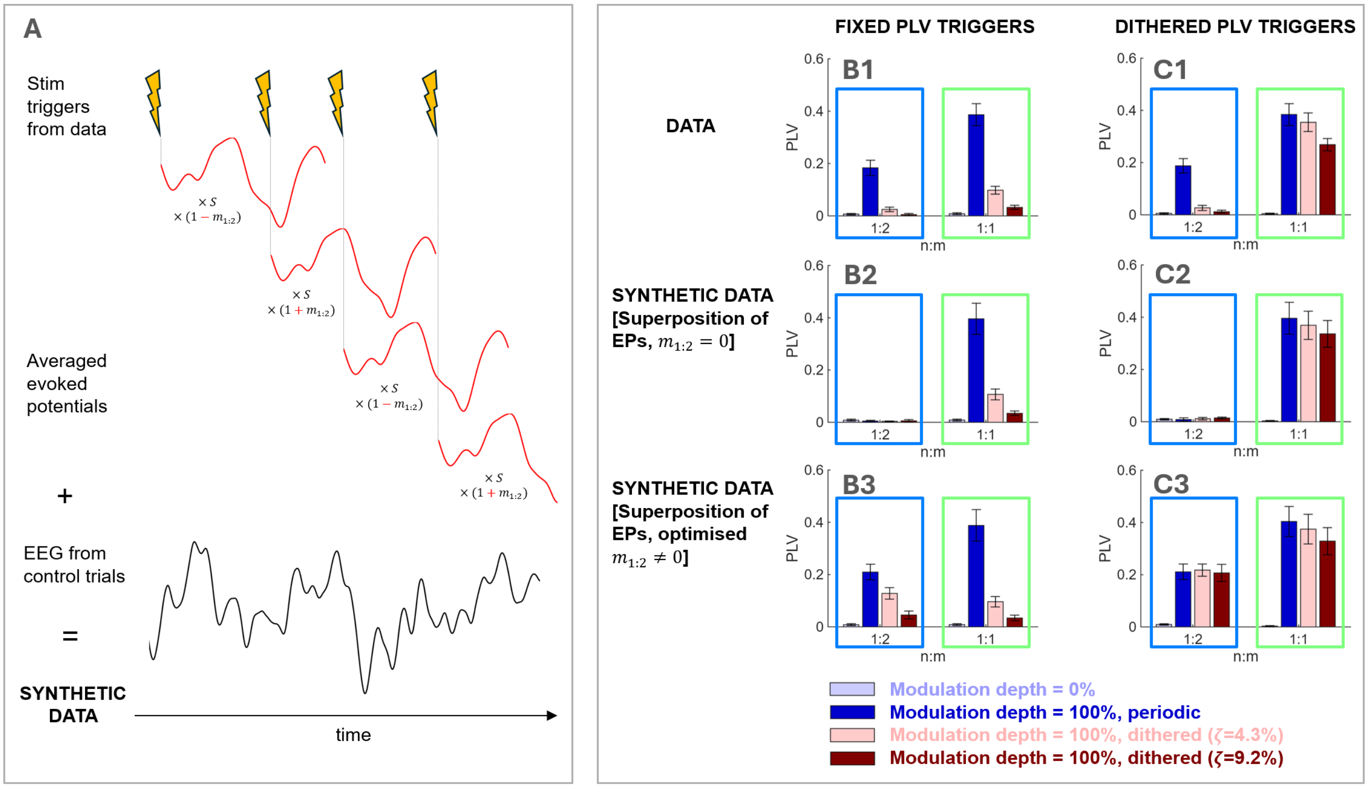


**Figure S.5: Half-harmonic responses to photic stimulation are inconsistent with the superposition of evoked potentials including frequency components at half the stimulation frequency A**: Schematic illustrating how synthetic data is generated according to the superposition of evoked potentials hypothesis, even when using averaged evoked potentials including frequency components at half the stimulation frequency. For each participant, the scale factor $S$ was determined to match the data $\text{PLV}_{1:1}$ for the periodic condition. In B3 and C3, the modulation factor $m_{1:2}$ was chosen to match the data $\text{PLV}_{1:2}$ for the periodic condition for each participant. **B-C**: PLV at the stimulation frequency (1:1) and its half-harmonic (1:2) at the group level, for fixed PLV triggers (B) and dithered PLV triggers (C). Comparison between the data and synthetic data generated according to the superposition of evoked potentials hypothesis (without and with modulation at the half-frequency). Panels B-C share the same legend, and error bars represent the standard error of the mean.
